# Atopic Biomarker Changes after Exposure to *Porphyromonas gingivalis* Lipopolysaccharide: A Small Experimental Study in Wistar Rat

**DOI:** 10.1101/2021.01.14.426656

**Authors:** Sindy Cornelia Nelwan, Ricardo Adrian Nugraha, Anang Endaryanto, Asti Meizarini, Udijanto Tedjosasongko, Seno Pradopo, Haryono Utomo

## Abstract

**Background:** IgE and IgG_4_ are implicated in atopic development and clinically utilized as major biomarkers. Atopic responses following certain pathogens such as *Porphyromonas gingivalis* is currently an area of interest for further research.

**Purpose:** The aim of this study is to measure the level of IgE, IgG_4_, and IgG_4_/IgE ratio periodically after exposure of periodontal pathogen *Porphyromonas gingivalis* (Pg) lipopolysaccharide (LPS).

**Methods:** We used 16 wistar rats (*Rattus norvegicus*) randomly subdivided into 4 groups, group 1 were injected by placebo, group 2 by LPS Pg 0.3 μg/mL,group 3 by LPS Pg 1 μg/mL, and group 4 by LPS Pg 3 μg/mL. Sera from both groups were taken from retro-orbital plexus before and after exposure.

**Results:** Level of IgE and IgG_4_ increased significantly following exposure of LPS Pg at day-4 and day-11. Greater increase of IgE rather than IgG_4_ contributes to rapid decline of IgG_4_/IgE ratio, detected in the peripheral blood at day-4 and day-11.

**Conclusion:** Modulation of atopic responses following exposure to LPS Pg is reflected by decrease in IgG_4_/IgE ratio that accompanies an increase of IgE.

**Clinical significance:** *Porphyromonas gingivalis*, a keystone pathogen during periodontal disease, may have a tendency to disrupt atopic biomarkers.

## INTRODUCTION

Oral cavity is the the habitat of numerous bacterias including *Porphyromonas gingivalis* (Pg). Pg is a gram negative, facultative anaerobic pathogen which is responsible in causing gingivitis or periodontitis (1). In low-income countries, gingivitis and periodontitis can affect up to 90% of the adult population (2). Rather than alveolar bone and ligament destruction, *Porphyromonas gingivalis* is believed to have some linked with the development of atopic responses in a susceptible host (3). Following Pg infection, hosts’ adaptive immune response (both cell-mediated and humoral-mediated) could induce systemic inflammatory reaction, not only just local destruction of tooth-supporting tissues (4–5). Although periodontal pathogens such as Pg play major role in the initiation of local and systemic inflammatory reaction (6), the host aberrant immune responses are more interesting subjects to be studied. Since humoral immune responses are stimulated following Pg infection, there might have been linked to the occurrence of atopy.

Despite long-standing research about hygiene hypothesis in several decade, there is an unequivocally-accepted fact that the prevalence of atopy increases greater among children who had periodontal pathogen colonization or infection (7). While endorsing these hygiene hypothesis approaches, there is an alternative hypothesis in which exposure to some periodontal pathogens will exclusively trigger an “Immunoglobulin-E skew” rather than declining it (8). Within the context of the hygiene hypothesis, the most essential microbial exposures need to be studied is biomolecular relationship between host antibody and regulatory T-cell with lipopolysaccharide (LPS), an endotoxin released by Pg to affect host immune reaction.

Hygiene hypothesis principles might not be able to answer all phenomenon of increasing incidence of atopy among children with poor oral hygiene (9). Some studies report a positive association between the colonization/infection of Pg with the development of allergic diseases (10–15), whereas some studies report no association (16–21). Due to lack of conclusive evidence about the association between Pg and allergic diseases (22), we try to measure the level of atopic biomarkers following Pg infection.

To the best of our knowledge, measuring IgG_4_ and IgE antibody may have a closer association to the atopic profiles, since IgG_4_ and IgE are released after activation of mature B cells following the modulation of IL-4 and IL-5 released by Th-2 cells during type I hypersensitivity (23). By looking at the alteration of IgG_4_ and IgE antibodies level after exposure to these selected components of Pg in an animal model, we hope to understand deeper about the biological mechanism of B-cell production antibodies patterns and humoral immune responses earlier before the clinical manifestation of atopy.

## MATERIAL AND METHODS

### Ethics approval

Animal experimental study were conducted under the approval of the Institutional Animal Research Ethics Committee of Universitas Airlangga (UNAIR), Surabaya, Indonesia (animal approval no:50/KKEPK.FKG/IV/2015) under the name of Sindy Cornelia Nelwan as the Principal Investigator. Study was carried out in strict accordance to internationally-accepted standards of the Guide for the Care and Use of Laboratory Animals of the National Institute of Health.

### Animals

The present study used 16 male Wistar rats (*Rattus novergicus*) between eight and ten weeks of age (average body weight 120-150 grams). The rats were housed in microisolator cages and maintained in a constant room temperature ranging from 22°C to 25°C, with a 12-h light/12-h dark cycle, under artificially controlled ventilation, with a relative humidity ranging from 50% to 60%. The rats were fed a standard balanced rodent diet (NUTRILAB CR-1®) and water were provided ad libitum.

### Experimental design and groups

The present study design was a pre-test post-test controlled group design using quantitative method. We extracted 16 male Wistar rats, randomized and then classified into 4 groups. Each group consists of 4 matched Wistar rats (age, weight, IgE and IgG_4_ baseline characteristic). Group 1 were given placebo. Group 2 were given lipopolysaccharide (LPS) of *Porphyromonas gingivalis* (Pg) at dose 0.3 μg/mL. Group 3 were given LPS Pg at dose 1 μg/mL. Group 4 were given LPS Pg at dose 3 μg/mL. The enrolled subjects will receive LPS by an intra-sulcular injection. Intra-sulcular injection has an advantages due to the its direct delivery of LPS to oral cavity in which the tip of needle is injected slowly at the crestal bone. Longitudinal quantitative measurement had been done by repeated checked of IgE level, IgG_4_ level, and IgG_4_/IgE ratio in both groups between day-0 (before treatment), day-4, and day-11. An average of 0.2 ml peripheral blood sera was obtained by Pasteur pipette from retro-orbital plexus, using a lateral approach.

### Level of IgG_4_ and IgE

Sample of the sera were collected and stored at −70°C (−94°F) at Institute of Tropical Diseases Universitas Airlangga (UNAIR). All sera were assessed by direct-sandwich enzyme-linked immunosorbent assay (ELISA) under the manufacturer’s (R&D System Europe Ltd, Abingdon, UK) protocol. Briefly, the sera were examined at the microtiter plates with mix supplied 25 ml of 3,3’,5,5’-tetramethylbenzidine to 1 ml of phosphate-citrate buffer plus perborate in a mildly acidic buffer (adjust pH 5.7). Level of IgG_4_ were detected using monoclonal antibody anti-IgG_4_, transferring it to microtiter plates, adding the supplied conjugate, adding blocking solution, diluting plasma sample (1:100,000), and washing between the steps. Level of IgE was detected using monoclonal antibody anti-IgE, similar following steps until diluting plasma sample (1:200). A minimum value of 0.01 pg/mL for IgE and 0.01 ng/mL for IgG_4_ were assigned for below the limit of detection.

### Statistical analysis

All measurements were performed at least three times. Results were presented as means ± standard errors (SEM). The assumption of the normality for the complete data was assessed by Shapiro-Wilk test. Test of homogeneity of variances was assessed by Levene Statistics. Statistical significance were examined by one-way ANOVA and repeated measure ANOVA using SPSS version 17.0 for Microsoft (IBM corp, Chicago, USA).

## RESULTS

### General characteristic investigations

Table 1 showed baseline characteristics of 16 Wistar rats (*Rattus norvegicus*). No significant differences had been found for mean age (*p*=0.774), body weight (*p*=0.700), baseline IgE (*p*=0.071), baseline IgG_4_ (*p*=0.770), and baseline IgG_4_/IgE ratio (*p*=0.053) among the four groups. (*Table* 1)

**Table 1.** General characteristics of study population

| Variable | Placebo | LPS Pg 0,3<br>ug/ml | LPS Pg 1<br>ug/ml | LPS Pg 3<br>ug/ml | *p value* |
| --- | --- | --- | --- | --- | --- |
| Age (weeks) | 8.87 ± 0.86 | 9.37 ± 0.95 | 8.75 ± 0.96 | 9.12 ± 0.63 | 0.774 |
| Weight (g) | 135.25 ± 9.54 | 135.25 ± 9.55 | 137.25 ± 7.93 | 138.00 ± 10.98 | 0.700 |
| Male (%) | 100 | 100 | 100 | 100 | - |
| IgE (pg/mL) | 5.69 ± 0.28 | 5.36 ± 0.19 | 10.27 ± 0.70 | 6.69 ± 0.61 | 0.071 |
| IgG<sub>4</sub> (ng/mL) | 10.74 ± 1.41 | 11.76 ± 0.85 | 11.91 ± 1.66 | 10.14 ± 1.44 | 0.770 |
| IgG<sub>4</sub>/IgE (x10<sup>3</sup>) | 1.87 ± 0.18 | 2.20 ± 0.13 | 1.19 ± 0.20 | 1.59 ± 0.34 | 0.053 |

**Notes:**
\* Data are presented as mean ± standard error of the mean (SEM)
\*\* One-way ANOVA for categorical variables; significant at $p < 0.05$
\*\*\* IgE, immune globulin E; IgG<sub>4</sub>, immune globulin G<sub>4</sub>; IgG<sub>4</sub>/IgE, ratio between averages IgG<sub>4</sub> levels divided by averages IgE levels.

### Comparison of serum IgE level between four groups

Prior to experiments (day-0), there was no difference of serum IgE level between the four groups (*p*>0.05). Meanwhile, four days following experiments, there was a significance difference of serum IgE level between both groups (*p*=0.006). At day-4, the highest average IgE level could be found in the group treated with LPS Pg 1 **μ**g/ml (17.00±1.69 pg/ml) and the lowest average IgE level could be found in the control group (5.31±0.76 pg/ml) respectively. Furthermore, ten days following experiments, there was also a significance difference of serum IgE level between both groups (*p*=0.047). At day-10, the highest average IgE level could be found in the group treated with LPS Pg 0.3 **μ**g/ml (180.34±10.42 pg/ml) and the lowest average IgE level could be found in control group (5.06±1.86 pg/ml) respectively. (*Table* 2)

**Table 2.** Comparisons of total serum IgE (pg/mL) between four groups (mean ± SEM)

| Group | n | IgE day 0 | IgE day 4 | IgE day 11 |
| --- | --- | --- | --- | --- |
| Placebo | 4 | 5.69 ± 0.28 | 5.31 ± 0.76 | 5.06 ± 1.86 |
| LPS Pg 0,3 ug/ml | 4 | 5.36 ± 0.19 | 9.26 ± 0.32 | 180.34 ± 10.42 |
| LPS Pg 1 ug/ml | 4 | 10.27 ± 0.70 | 17.00 ± 1.69 | 112.90 ± 3.87 |
| LPS Pg 3 ug/ml | 4 | 6.69 ± 0.61 | 14.79 ± 0.86 | 102.01 ± 11.04 |
| F statistic | / | 20.733 | 26.171 | 83.758 |
| *p* value | / | 0.071 | 0.006 | 0.047 |

\* measured by one-way ANOVA (df1=3, df2=12, f table 3.490; significant at $p < 0.05$ )

### Comparison of serum IgG_4_ level between four groups

Prior to experiments (day-0), there was no difference of serum IgG_4_ level between the four groups (*p*>0.05). Meanwhile, four days following experiments, there was a significance difference of serum IgG_4_ level between both groups (*p*=0.008). At day-4, the highest average IgG_4_ level could be found in the group treated with LPS Pg 3 **μ**g/ml (23.86±1.59 ng/ml) and the lowest average IgG_4_ level could be found in the control group (8.34±0.58 ng/ml) respectively. Furthermore, ten days following experiments, there was a greater difference of serum IgG_4_ level between both groups (*p*=0.005). At day-10, the highest average IgG_4_ level could be found in the group treated with LPS Pg 3 **μ**g/ml (63.74±4.74 ng/ml) and the lowest average IgG_4_ level could be found in control group (13.91±0.99 ng/ml) respectively. (*Table* 3)

**Table 3.**
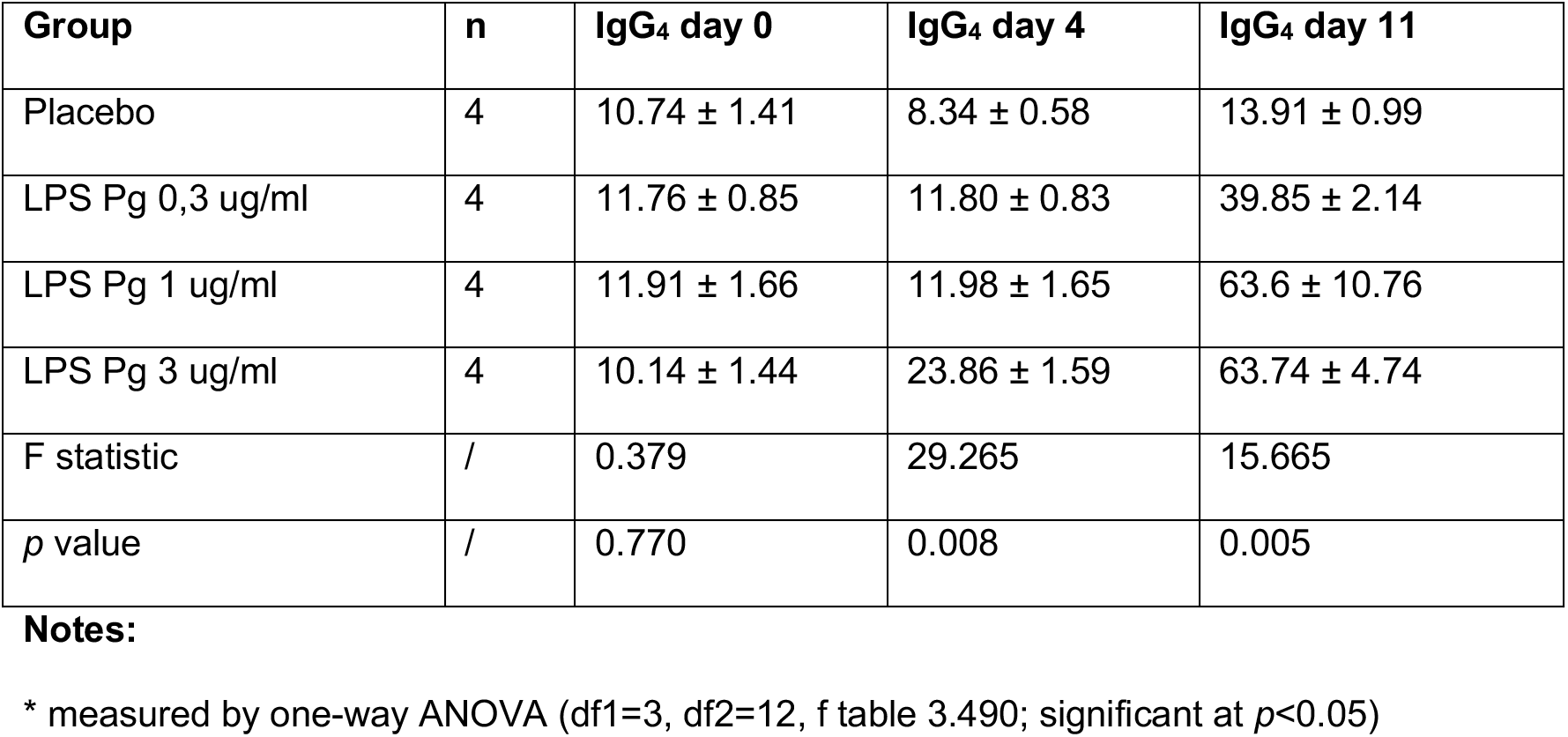
Comparisons of total serum IgG_4_ (ng/mL) between four groups (mean ± SEM)

### Ratio of IgG_4_/IgE antibodies between four groups

The average IgG_4_/IgE ratio for the control group at day-0, day-4, and day-11 was 1.87×10^3^, 1.72×10^3^, and 3.62×10^3^. In the low-dose LPS group, the average IgG_4_/IgE ratio was 2.20×10^3^, 1.27×10^3^, and 0.22×10^3^. In the medium-dose LPS group, the average IgG_4_/IgE ratio was 1.19×10^3^, 0.69×10^3^, and 0.56×10^3^. In the high-dose LPS group, the average IgG_4_/IgE ratio was 1.59×10^3^, 1.64×10^3^, and 0.65×10^3^. All groups exhibited significant differences of IgG_4_/IgE ratios, except at day-0. The highest IgG_4_/IgE ratio at day-4 and day-10 could be found in control group. The lowest IgG_4_/IgE ratio at day-4 could be found in LPS Pg 1 **μ**g/ml group, whilst the lowest ratio at day-11 could be found in LPS Pg 0.3 **μ**g/ml respectively. (*Table* 4)

**Table 4.**
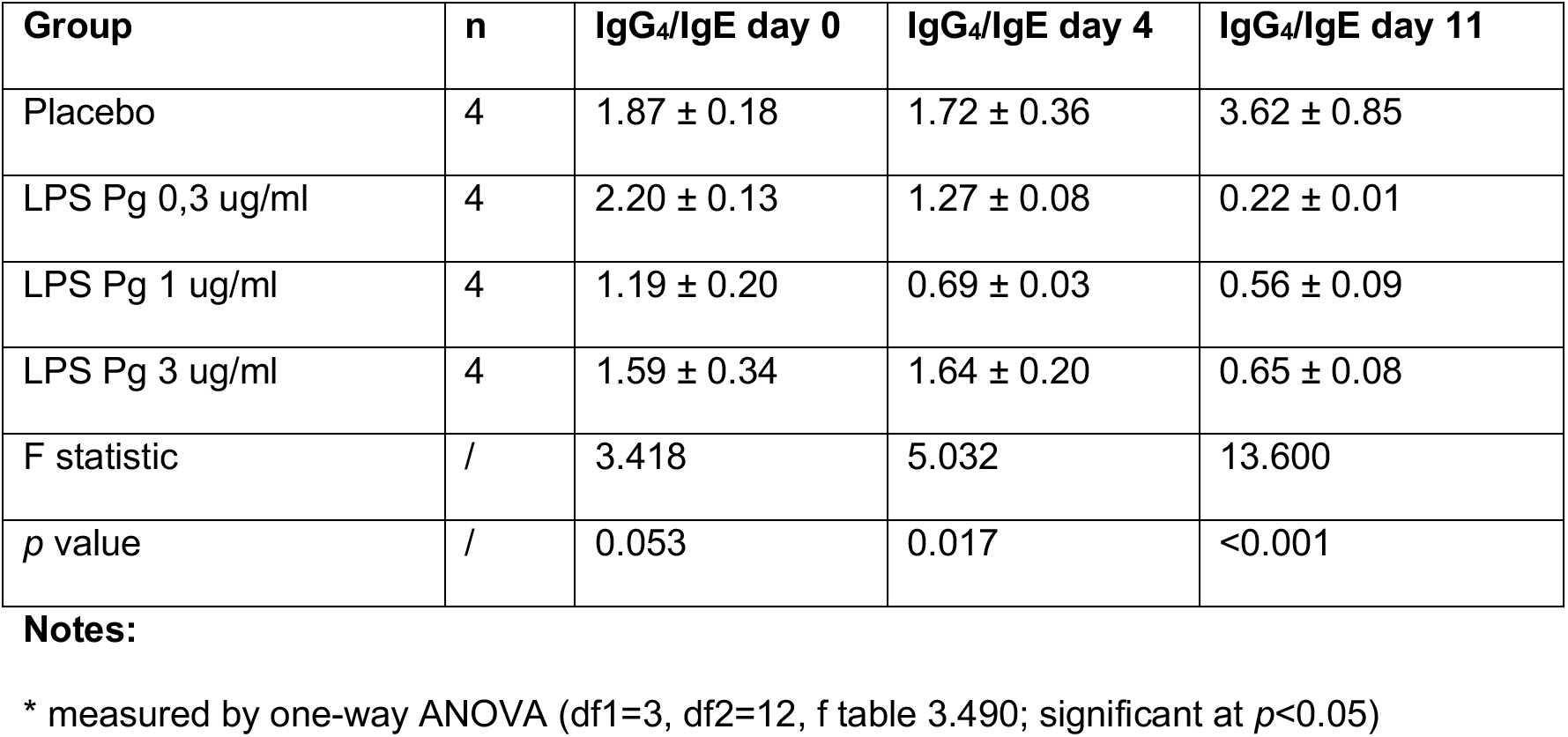
Comparisons of IgG_4_/IgE ratio (x10^3^) between four groups (mean ± SEM)

| Group | n | IgG<sub>4</sub>/IgE day 0 | IgG<sub>4</sub>/IgE day 4 | IgG<sub>4</sub>/IgE day 11 |
| --- | --- | --- | --- | --- |
| Placebo | 4 | 1.87 ± 0.18 | 1.72 ± 0.36 | 3.62 ± 0.85 |
| LPS Pg 0,3 ug/ml | 4 | 2.20 ± 0.13 | 1.27 ± 0.08 | 0.22 ± 0.01 |
| LPS Pg 1 ug/ml | 4 | 1.19 ± 0.20 | 0.69 ± 0.03 | 0.56 ± 0.09 |
| LPS Pg 3 ug/ml | 4 | 1.59 ± 0.34 | 1.64 ± 0.20 | 0.65 ± 0.08 |
| F statistic | / | 3.418 | 5.032 | 13.600 |
| *p* value | / | 0.053 | 0.017 | <0.001 |

\* measured by one-way ANOVA (df1=3, df2=12, f table 3.490; significant at *p*<0.05)

### Subgroup analysis in Wistar rats treated with of 0.3 μg/ml LPS Pg

Level of IgE were increased dramatically from day-0 to day-11 after experiments (5.36±0.19 pg/ml to 180.34±10.42 pg/ml; *p*=0.011). Level of IgG_4_ were also increase significantly from day-0 to day-11 after experiments (11.76±0.85 ng/ml to 39.85±2.14 ng/ml; *p*=0.006). On the other hand, IgG_4_/IgE ratio were decreased following experiments (2.20±0.13×10^3^ to 0.22±0.01×10^3^; *p*=0.014). (*Table* 5)

**Table 5.** Comparisons of total serum IgE, IgG_4_, and IgG_4_/IgE ratio before and after treatment in the group with exposure of 0.3 μg/mL LPS Pg (mean ± SEM)

| Time-point | n | IgE (pg/mL) | IgG<sub>4</sub> (ng/mL) | IgG<sub>4</sub>/IgE (x10<sup>3</sup>) |
| --- | --- | --- | --- | --- |
| Day-0 Before treatment | 4 | 5.36 ± 0.19 | 11.76 ± 0.85 | 2.20 ± 0.13 |
| Day-4 After treatment | 4 | 9.26 ± 0.32 | 11.80 ± 0.83 | 1.27 ± 0.08 |
| Day-11 After treatment | 4 | 180.34 ± 10.42 | 39.85 ± 2.14 | 0.22 ± 0.01 |
| F statistic | / | 93.924 | 178.233 | 71.969 |
| p value | / | 0.011 | 0.006 | 0.014 |

\* measured by repeated measure ANOVA
\*\* df times=2, df error=14, f table 3.739; significant at $p < 0.05$

### Subgroup analysis in Wistar rats treated with 1 μg/ml LPS Pg

Level of IgE were raised dramatically from day-0 to day-11 after experiments (10.27±0.70 pg/ml to 112.90±3.87 pg/ml; *p*=0.003). Level of IgG_4_ were also increase significantly from day-0 to day-11 after experiments (11.91±1.66 ng/ml to 63.6±10.76 ng/ml; *p*=0.027). On the other hand, IgG_4_/IgE ratio were declined following experiments (1.19±0.20×10^3^ to 0.56±0.09×10^3^; *p*=0.362). (*Table* 6)

**Table 6.** Comparisons of total serum IgE, IgG_4_, and IgG_4_/IgE ratio before and after treatment in the group with exposure of 1 μg/mL LPS Pg (mean ± SEM)

| Time-point | n | IgE (pg/mL) | IgG<sub>4</sub> (ng/mL) | IgG<sub>4</sub>/IgE (x10<sup>3</sup>) |
| --- | --- | --- | --- | --- |
| Day-0 Before treatment | 4 | 10.27 ± 0.70 | 11.91 ± 1.66 | 1.19 ± 0.20 |
| Day-4 After treatment | 4 | 17.00 ± 1.69 | 11.98 ± 1.65 | 0.69 ± 0.03 |
| Day-11 After treatment | 4 | 112.90 ± 3.87 | 63.6 ± 10.76 | 0.56 ± 0.09 |
| F statistic | / | 294.526 | 35.437 | 1.760 |
| p value | / | 0.003 | 0.027 | 0.362 |

\* measured by repeated measure ANOVA
\*\* df times=2, df error=14, f table 3.739; significant at $p < 0.05$

### Subgroup analysis in Wistar rats treated with of 3 μg/ml LPS Pg

Level of IgE were raised dramatically from day-0 to day-11 after experiments (6.69±0.61 pg/ml to 102.01±11.04 pg/ml; *p*=0.009). Level of IgG_4_ were also increase significantly from day-0 to day-11 after experiments (10.14±1.44 ng/ml to 63.74±4.74 ng/ml; *p*=0.029). On the other hand, IgG_4_/IgE ratio were declined following experiments (1.59±0.34×10^3^ to 0.65±0.08×10^3^; *p*=0.113). (*Table* 7)

**Table 7.** Comparisons of total serum IgE, IgG_4_, and IgG_4_/IgE ratio before and after treatment in the group with exposure of 3 μg/mL LPS Pg (mean ± SEM)

| Time-point | n | IgE (pg/mL) | IgG<sub>4</sub> (ng/mL) | IgG<sub>4</sub>/IgE (x10<sup>3</sup>) |
| --- | --- | --- | --- | --- |
| Day-0 Before treatment | 4 | 6.69 ± 0.61 | 10.14 ± 1.44 | 1.59 ± 0.34 |
| Day-4 After treatment | 4 | 14.79 ± 0.86 | 23.86 ± 1.59 | 1.64 ± 0.20 |
| Day-11 After treatment | 4 | 102.01 ± 11.04 | 63.74 ± 4.74 | 0.65 ± 0.08 |
| F statistic | / | 111.386 | 32.929 | 7.841 |
| p value | / | 0.009 | 0.029 | 0.113 |

**Notes:** \* measured by repeated measure ANOVA \*\* df times=2, df error=14, f table 3.739; significant at $p < 0.05$ .

## DISCUSSION

Several mechanisms have been suggested to alter to atopic inflammatory responses following LPS Pg infection. One of the mechanisms proven in this study is an elevation of IgE antibody and declining of IgG_4_/IgE ratio (24). Other mechanisms which are indirectly proven are shifting of Th-1 to become Th-2 cells along with their cytokines pathway. Down-regulation of the Th-1 cell associated to depress of cell-mediated immune response and stimulate humoral immune response, thus pathogens are able to evade the immune clearance (25).

*Porphyromonas gingivalis* possess very sophisticated defense mechanisms against host immune responses. These pathogens produce capsules containing long chain LPS which is designed effectively to counter MAC. These long chain LPS can also downgrade cell-mediated immunity by shifting Th-1 into Th-2 which less dangerous to pathogens (26). LPS may have an essential role in switching cell-mediated to humoral-mediated immune responses (27). LPS Pg antigen is processed and presented on its surface again with MHC-II molecule. Recent studies suggest an activation of alternative complement pathway, disruption of classical complement pathway, modulation of antigen presenting cells, and downregulation of anti-inflammatory cytokines are responsible for the Th2-skewed immune response following exposure to LPS Pg. Predominance shifting from Th-1 into Th-2 occurs in several extra-lymphoid tissues; the ideal site for *Porphyromonas gingivalis* is the oral cavity (28).

Interleukin-4 (IL-4), which is produced by naive T cells, acts as autocrine manner known to be responsible for the differentiation and activation of Th-2 phenotype (29). Guo et al (2014) shows upon the occurrence and development of allergic diseases, there is a complex pathobiology which results an imbalance of Th-1/Th-2 (30). In an atopic disease such as bronchial asthma or urticaria, naive T cell can differentiate into Th-2 under the spesific circumstance by IL-4-induced STAT6 and GATA-3 transcription factors (30). Th-2 predominant immune response will automatically stimulate plasma cell to release IgE and IgG_4_ (31). Upon re-exposure of antigen or allergen, binding of the allergen to IgE orchestrates the adaptive immune system to initiate rapid sensitization. Frequent sensitization is a major risk factor for the development of allergic diseases such as urticaria, bronchial asthma, hay fever or atopic dermatitis/ eczema (32).

Our previous study used whole-cell body of *Porphyromonas gingivalis* to study different molecular responses in Wistar rats. Our first project studied the association between periodontal pathogen and host innate immunity. Exposure to *Porphyromonas gingivalis* had been shown to stimulate level of TLR2 and depress level of TLR4 (33). Our findings might indicate that several bacterial properties can turn-off host innate immunity and host inflammatory response. Our second project studied the association between periodontal pathogen and host adaptive immunity. We summarized that high dose CFU of Pg stimulates fold increase of Th-2 cytokines (IL-4, IL-5 and IL-13) and decrease of Th-1 cytokines (IFN-γ and IL-17) (34). These were the cornerstone to continue our project in studying LPS as the most important component of these bacteria.

At this moment, both total IgE or spesific IgE antibodies have little diagnostic value in the occurence of allergic manifestation. Even total or spesific IgE is increasing, yet the manifestation of allergy doesn’t usually develop, since IgG_4_ level is also increase as a counter-regulator (35). It means that even human or rat become susceptible to atopic due to the increasing level of IgE, body mechanism is able to provide protection, with increased IgG_4_ as a counter response to prevent manifestation of allergic diseases and immediate hypersensitivity. Thus, exposure of LPS Pg will develop chance of atopic and hypersensitivity markers, but manifestation of allergic reaction is a complex pattern (36). IgG_4_/IgE ratio has closer accuracy to detect any alteration of atopic inflammatory pathway. Increase level of IgE which isn’t accompanied by IgG_4_ can be seen in patient with urticaria or atopic dermatitis (37). IgE-switched B cells are much more likely to differentiate into plasma cells, whereas IgG_4_-switched B cells are less likely to differentiate (38). This reason would explain why IgE antibody is the most dominant antibodies in the development of atopic inflammatory pathway, whereas IgG_4_ antibodies become prominent later during chronic non-atopic stimulation (39). According to this reason, IgG_4_/IgE ratio may predict atopic responses more accurately than total or specific IgE level.

### Limitations and Strength

Several limitations should be highlighted. First, this study had limitations with regard to very small number of samples which can increase the likelihood of error and imprecision. Second, results from animal model often do not translate into replications in human model (40). IgE antibody responses in Wistar rats are typically transient, whereas the atopic IgE response in human persists for many years (41). Other crucial difference is IgG_4_/IgE ratio, which is usually much higher in the Wistar rats than human (42–43). These factors may have an impact on the interpretation of our results. Thus, the findings should be interpreted within the context of this study and its limitations. The strengths of the study were its high statistical power and the homogeneity of each group to enable comparison between groups and periods.

### Conclusion

Several experiments in rats indicate that exposure to LPS Pg may have a tendency to increase level of IgE and IgG_4_. On the contrary, declining of IgG_4_/IgE ratio following exposure to LPS Pg suggest the potential role of LPS Pg for isotype switching from IgG_4_ to IgE. The results of the present study favor indirect isotype switching route for most IgE as secondary responses from LPS Pg infection that leads to systemic atopic inflammatory pathway.

**Figure 1.**
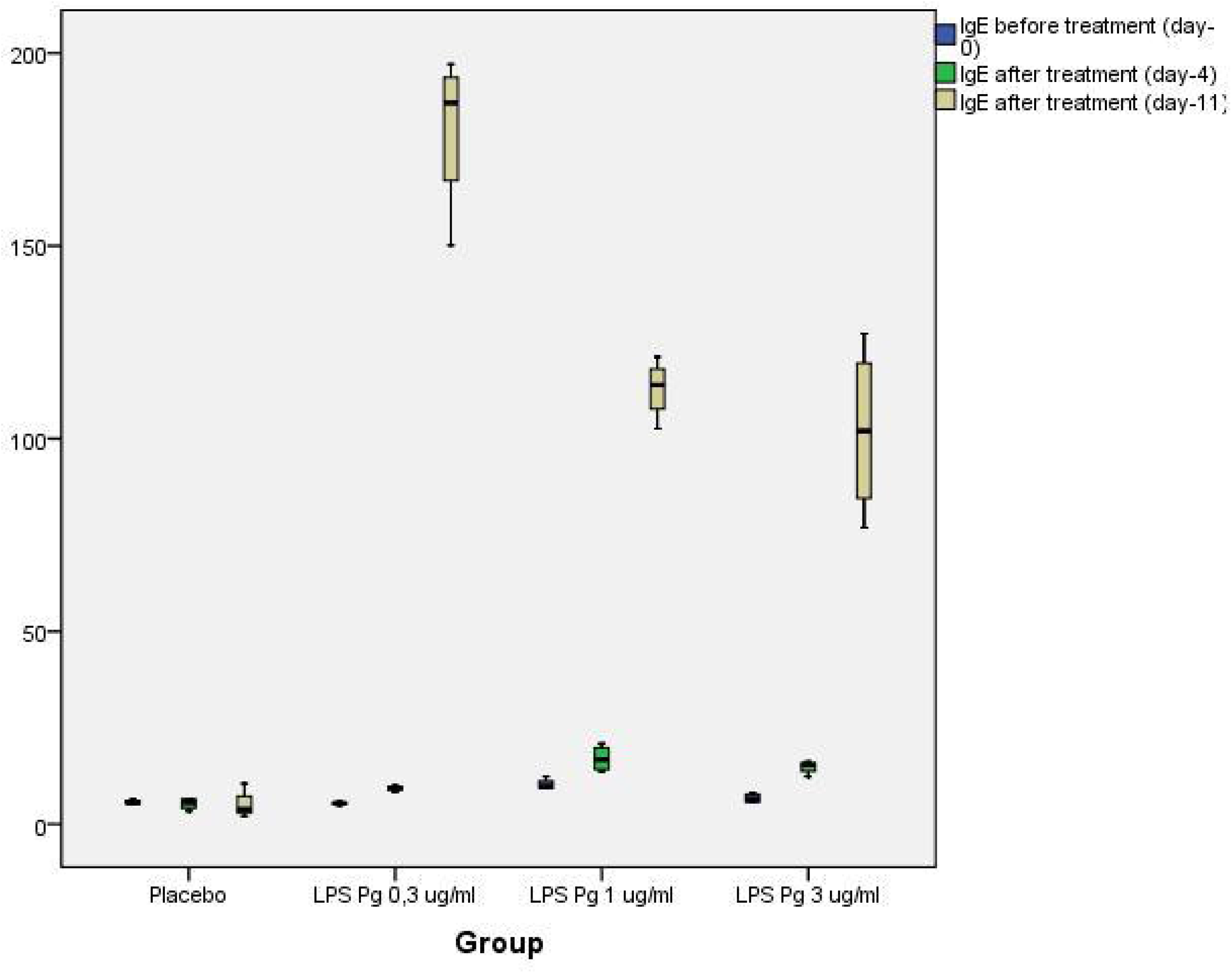
Longitudinal observation of serum IgE levels following exposure to LPS Pg.

**Figure 2.**
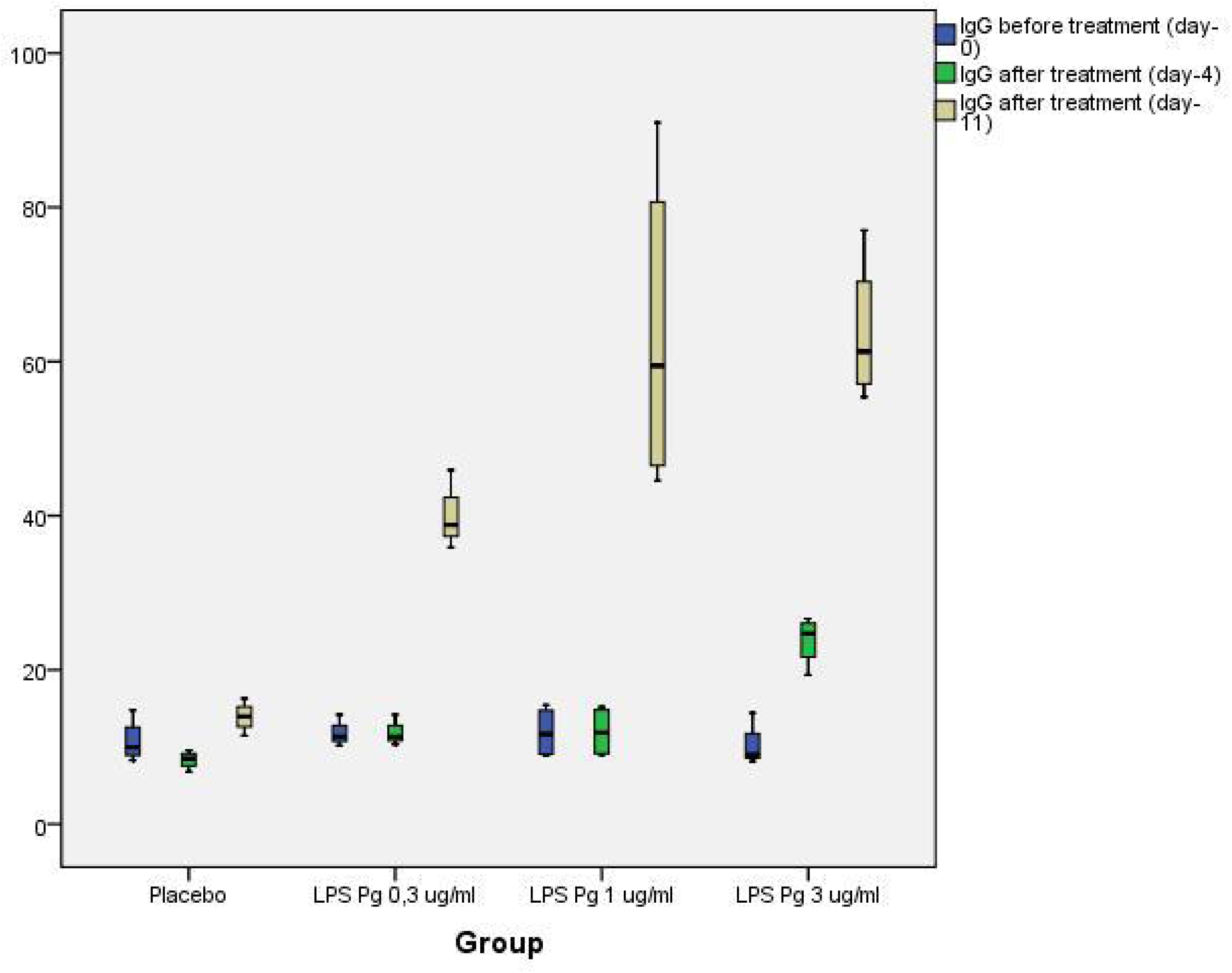
Longitudinal observation of serum IgG_4_ levels following exposure to LPS Pg.

**Figure 3.**
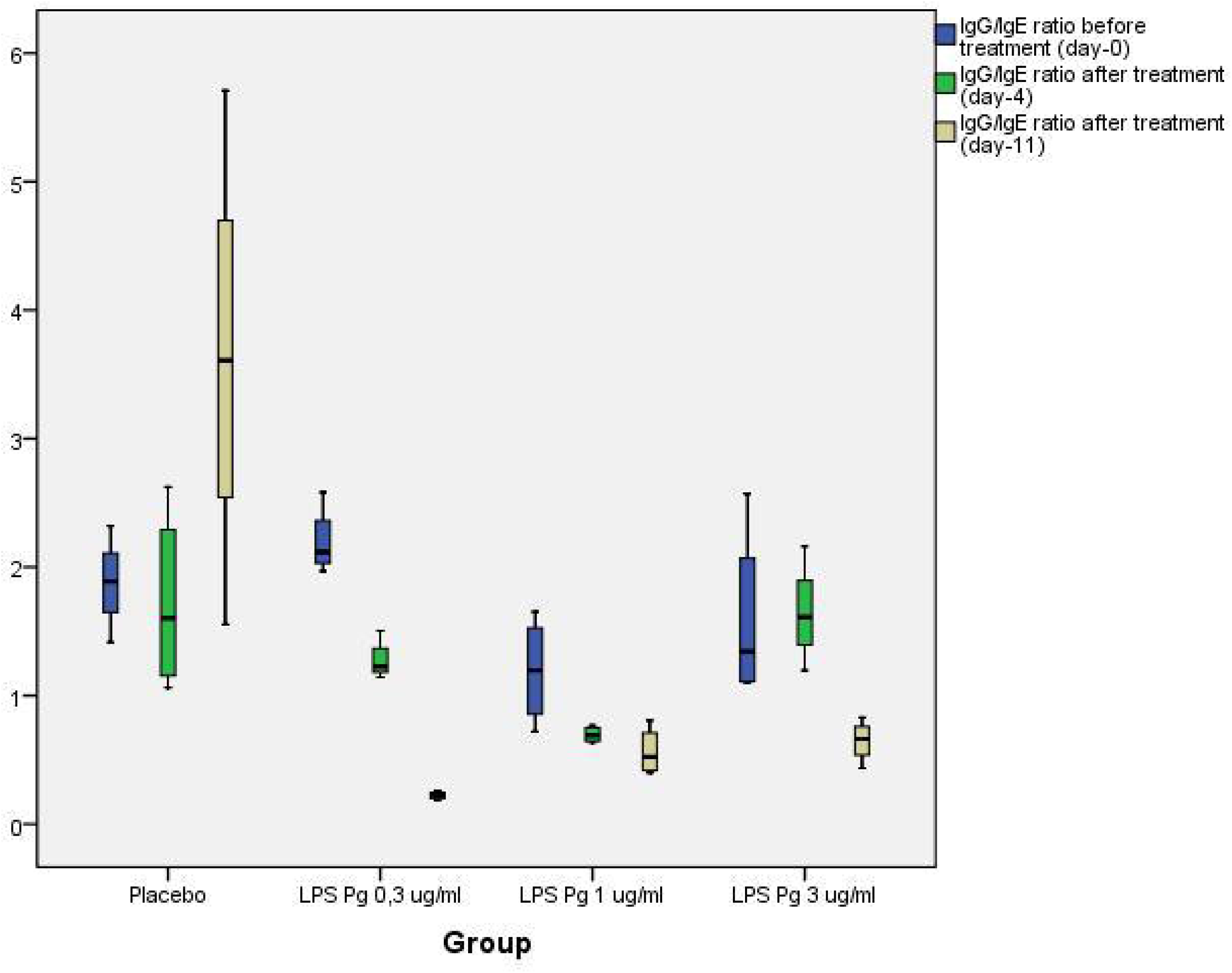
Comparisons of IgG_4_/IgE ratio between four groups (mean ± SEM).

## ACKNOWLEDGMENTS

This project is fully supported by Darmawan Setijanto as Dean of Universitas Airlangga College of Dentistry and Coen Pramono as Director of Airlangga Oral and Dental Hospital. We would like to acknowledge Harjanto from Physiology Department and Indrawati Retno from Biochemistry Department for providing material and reagent generation. We also thank all lecturers and research assisstants from Universitas Airlangga, who are willing to help in the technical aspect.

## AUTHOR CONTRIBUTIONS

Conceptualization: Nelwan SC, Endaryanto A.

Project administration and funding acquisition: Nelwan SC, Endaryanto A, Utomo H.

Data curation and formal analysis: Nugraha RA.

Investigation: Nelwan SC, Nugraha RA.

Methodology: Nelwan SC, Nugraha RA.

Resources and Software: Nelwan SC, Meizarini A, Utomo H.

Supervision and validation: Tedjosasongko U, Pradopo S.

Writing - original draft: Nugraha RA.

Writing - review & editing: Tedjosasongko U, Utomo H.

## AVAILABILITY OF DATA AND MATERIALS

The datasets used and/or analyzed during the current study are available online at Universitas Airlangga Repositroy (http://repository.unair.ac.id/id/eprint/63285) for research purposes.

## CONSENT FOR PUBLICATIONS

Not applicable (public data).

## ABBREVIATIONS

ANOVA: analysis of variant
BCR: B cell receptor
CD-40: Cluster Differentiation-40
CFU: colony-forming unit
CSR: class-switch recombination
df: degree of freedom
ELISA: enzyme-linked immunosorbent assay
FDA: Food and Drug Administration
HDM: house dust mite
IACUC: Institutional Animal Care and Use Committee
IFN-γ: gamma-Interferon
IgE: immunoglobulin-E
IgG_4_: immunoglobulin-G4
IL: interleukin
LLPC: long-lived plasma cell
LPS: lipopolysaccharide
mAb: monoclonal antibody
MAC: membrane attack complex
MHC: major histocompatibility complex
NIH: National Institutes of Health
NGS: next-generation sequencing
NK: natural killer cells
OIT: oral immunotherapy
PAMP: pathogen-associated molecular patterns
PCR: Polymerase Chain Reaction
Pg: *Porphyromonas gingivalis*
SD: standard deviation
SEM: standard error of the mean
SPSS: Statistical Package for the Social Sciences
Th-1: type-1 helper T-cells
Th-2: type-2 helper T-cells
TNF-α: Tumor Necrosis Factor-α

